# Interleukin-33 from oligodendrocytes sustains effector differentiation of tissue-resident CD8+ T cells and is a druggable target in CNS autoimmune disease

**DOI:** 10.1101/2024.07.09.602661

**Authors:** Nicolas Fonta, Nicolas Page, Bogna Klimek, Margot Piccinno, Giovanni Di Liberto, Sylvain Lemeille, Mario Kreutzfeldt, Anna Lena Kastner, Yusuf I. Ertuna, Ilena Vincenti, Ingrid Wagner, Daniel D. Pinschewer, Doron Merkler

## Abstract

In chronic inflammatory disorders of the central nervous system (CNS), tissue-resident self-reactive T cells perpetuate disease. The specific tissue factors governing the persistence and continuous differentiation of these cells remain undefined but could represent attractive therapeutic targets.

In a model of chronic CNS autoimmunity, we find that oligodendrocyte-derived interleukin-33 (IL-33), an alarmin, is key for locally regulating the pathogenicity of self-reactive CD8+ T cells. The selective ablation of IL-33 from neo-self-antigen-expressing oligodendrocytes mitigates CNS disease. In this context, fewer self-reactive CD8+ T cells persist in the inflamed CNS, and the remaining cells are largely locked into a stem-like precursor program, failing to form TCF-1^low^ effector progeny. Importantly, interventional IL-33 blockade by locally administered somatic gene therapy reduces T cell infiltrates and improves the disease course.

Our study identifies oligodendrocyte-derived IL-33 as a druggable tissue factor regulating the differentiation and survival of self-reactive CD8+ T cell in the inflamed CNS. This finding introduces tissue factors as a novel category of immune targets for treating chronic CNS autoimmune diseases.

## Introduction

CD8+ T cells mediate pathogen elimination and tumor immunosurveillance, however, when aberrantly activated, they can contribute to the pathogenesis of a wide spectrum of chronic autoimmune diseases (Collier et al., 2021). In multiple sclerosis (MS), a chronic autoimmune disease of the central nervous system (CNS), CD8+ T cells constitute the majority of lymphocytes found in active-demyelinating CNS lesions, and they are considered critical effectors of tissue damage (Friese and Fugger, 2005). Importantly, circulating CD8+ effector memory (T_EM_) as well as tissue-resident memory T cells (T_RM_) are both important for perpetuating inflammation within the affected organ (Clark, 2015). While T_EM_ are in equilibrium with the circulation, T_RM_ cause tissue damage and sustain localized inflammation in a compartmentalized manner (Frieser et al., 2022; Vincenti et al., 2022).

Under conditions of prolonged antigen exposure such as in chronic viral infections and cancer, tissue infiltrating CD8+ T cells can undergo distinct differentiation reprogramming, which may lead to a progressive alteration of T cell effector function, often referred to as exhaustion (McLane et al., 2019). While thought to prevent excessive CD8+ T cell activity and tissue damage, this process of adaptation can further contribute to the persistence of autoreactive immune responses within the affected organ (Grebinoski et al., 2022). Among various factors that are critical for the persistence and the potential to sustain inflammation of autoreactive CD8+ T cells, the T cell-specific transcription factor T cell factor 1 (TCF-1, encoded by *Tcf7*) has garnered attention (Gearty et al., 2022). In the context of acute viral infection, CD8+ T cell-intrinsic TCF-1 is crucial for the development and preservation of functional memory CD8+ T cells (Zhou et al., 2010). Likewise, TCF-1^high^ CD8+ T cells retain stem-like properties when chronically stimulated, as has been demonstrated by their sustained proliferative potential, self-renewing capacity, and the ability to generate progeny cells that differentiate into terminal effector cells (Utzschneider et al., 2016; Wu et al., 2016). Although extrinsic factors such as the inflammatory cytokine interleukin-12 (IL-12) have been identified as potent regulators of the stemness function of CD8+ T cells (Danilo et al., 2018), the local cues within the target tissue that govern the longevity and the destructive potential of autoreactive CD8+ T cells are not well understood.

IL-33, an IL-1 family member, has been shown to act as a damage-associated molecular pattern (DAMP) or “alarmin” regulating a broad range of inflammatory processes (Liew et al., 2016). It is released upon tissue damage in the context of various pathological conditions such as in viral infections (Kallert et al., 2017; Bonilla et al., 2012), asthma (Préfontaine et al., 2009), allergy (Chan et al., 2019), and sepsis (Alves-Filho et al., 2010). The cytokine is sensed by immune cells (Cayrol and Girard, 2014) expressing the IL-33 receptor ST2 (IL-1RL1), amongst them Th1- and Th2-differentiated CD4+ (Baumann et al., 2015; Löhning et al., 1998) and CD8+ T cells (Bonilla et al., 2012). With regard to CD8+ T cell responses, IL-33 enhances the antiviral functionality of effector CD8+ T cells in the context of acute and latent infection as well as upon vaccination (McLaren et al., 2019; Kallert et al., 2017; Villarreal et al., 2014; Bonilla et al., 2012). Moreover, it preserves and promotes the stemness of memory-like CD8+ T cells in systemic viremic infection (Marx et al., 2023). Emerging evidence suggests further that IL-33 serves immunomodulatory functions in the CNS, where it is expressed at high levels (Fairlie-Clarke et al., 2018). The primary sources of IL-33 in the CNS are astrocytes during development (Vainchtein et al., 2018) and oligodendrocytes in adulthood (Gadani et al., 2015). In neurological disorders such as in multiple sclerosis (MS), it has been shown that IL-33 expression in both the peripheral nervous system and in CNS tissue is increased above steady-state levels in healthy controls (Jafarzadeh et al., 2016; Allan et al., 2016). This finding suggested that IL-33 may play a role in the pathophysiology of the disease. Yet, the exact role of IL-33 in chronic neuroinflammatory CNS disease remains incompletely understood, with reportedly pro-inflammatory but also anti-inflammatory effects (De la Fuente et al., 2015; Liew et al., 2016). In particular, the impact of glial cell-derived IL-33 on autoreactive CNS-infiltrating CD8+ T cells has remained unexplored.

Exploiting a CD8+ T cell-driven experimental autoimmune mouse model for chronic CNS inflammation, we investigated the impact of oligodendrocyte-derived IL-33 on CNS-infiltrating self-reactive CD8+ T cells, their transcriptional reprogramming and differentiation in the tissue environment and, consequently, tissue-destructive effects. Our investigations reveal a previously unrecognized role of the alarmin IL-33 as an oligodendrocyte-derived tissue factor sustaining the pathogenic responses of CNS-resident self-reactive CD8+ T cells, offering new avenues for therapeutic intervention.

## Results

### Oligodendrocytes-derived IL-33 enhances CD8+ T cell-mediated CNS autoimmune disease and tissue damage.

We set out to revisit the cellular sources of IL-33 and its expression levels in the CNS of healthy adult mice. Immunofluorescent co-labeling of IL-33 and OLIG2 – a transcription factor (TF) primarily expressed by cells of the oligodendrocyte lineage from precursor to maturity-confirmed a substantial overlap between these markers (Fig. 1A). In line with earlier findings suggesting oligodendrocytes as the primary cellular source of IL-33 in the adult CNS (Gadani et al., 2015), 64.17 ± 7.63% and 70.49 ± 8.44% of IL-33-expressing cells in brain and spinal cord respectively were from the oligodendrocyte lineage (OLIG2+ of IL-33+)(Fig. 1B). Hence, we generated a conditional knockout mouse strain lacking IL-33 selectively in mature oligodendrocytes (IL-33 cKO mice, Fig. 1A). For this, we crossbred mice engineered to express Cre recombinase as a knock-in under control of the myelin oligodendrocyte glycoprotein (MOG) promoter (MOGiCre, (Hövelmeyer et al., 2005)) to mice carrying a floxed IL-33 gene (Chen et al., 2015). When compared to their wildtype (WT) counterpart, IL-33 cKO mice exhibited a profound decrease in the proportion of IL-33+ cells among OLIG2+ oligodendrocytes (Fig. 1C), as intended.

**Figure 1:**
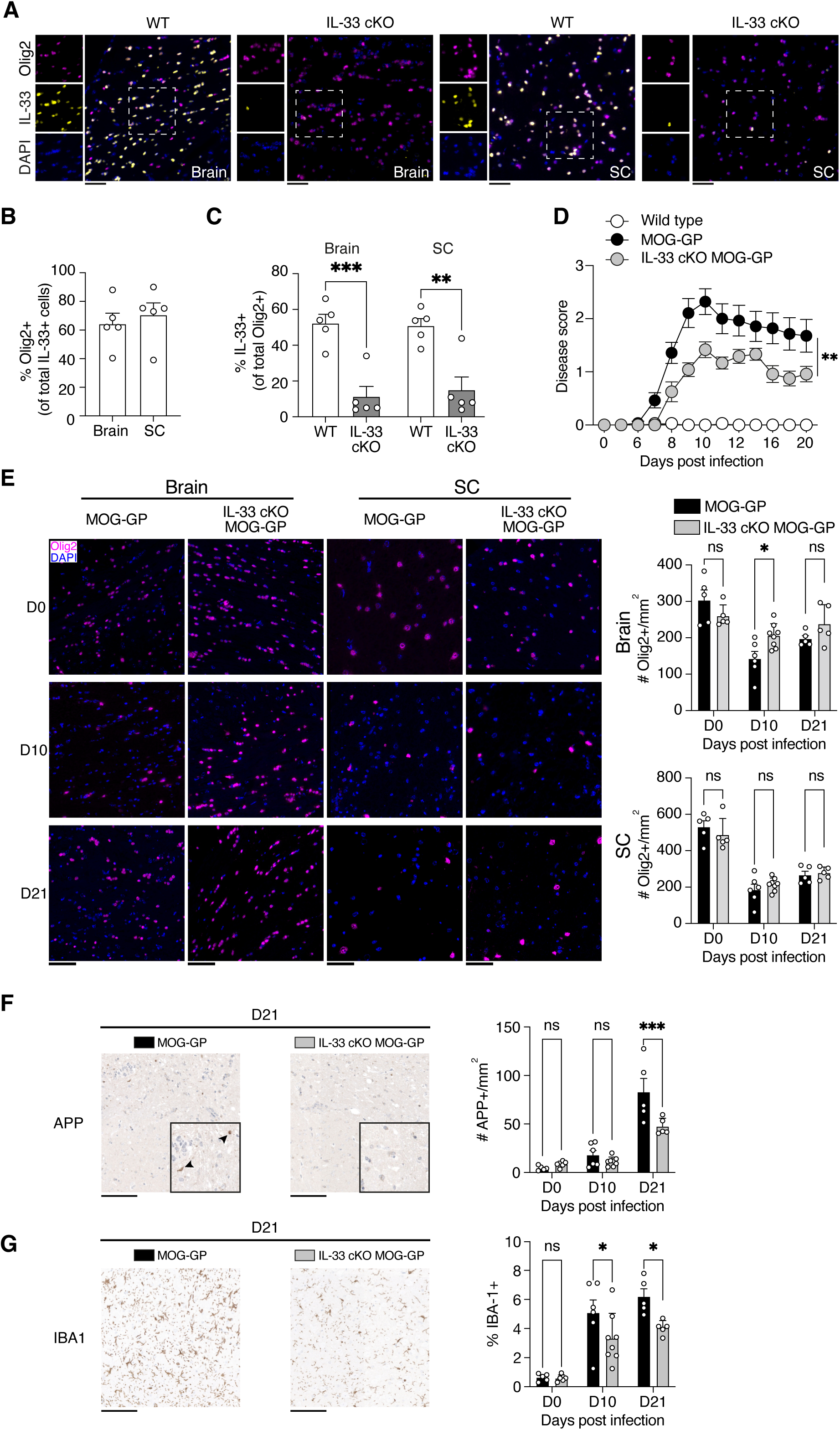
Oligodendrocytes-derived IL-33 enhances CD8+ T cell-mediated CNS autoimmune disease and tissue damage. **(A)** Brain and spinal cord (SC) sections of WT and IL-33 cKO mice stained for Olig2, IL-33, and Dapi. Scale bars, 50 μm **(B)** Percentage of Olig2+ cells among total IL-33+ cells in WT brain and spinal cord tissues. **(C)** Percentage of IL-33+ cells among total Olig2+ cells in WT and IL-33 cKO brain and spinal cord tissues. **(D-G)** WT, MOG-GP, and IL-33 cKO MOG-GP mice were infected i.v. with 10^4^ PFU rLCMV-GP33. **(D)** EAE disease course (n = 10 WT mice; n= 14 MOG-GP mice; n = 12 IL-33 cKO MOG-GP), clinical scores are expressed as mean ± SEM. **(E)** (Left) Representative immunostainings for Olig2 and Dapi on brain (Right) and spinal cord (Left) sections of MOG-GP and IL-33 cKO MOG-GP at indicated timepoint after rLCMV-GP33 infection. Scale bars, 50 μm. (Right) Number of Olig2+ cells/mm^2^ in brain (up) and spinal cord (down) tissues of indicated groups. **(F)** (Right) Representative immunostainings for APP in brain sections at day 21 in indicated groups. Arrowheads indicate axonal damage. Scale bars, 100 μm. (Left) Histological enumeration of APP+ cells/mm^2^ in brain at 0-, 8- and 21 days post-infection. **(G)** (Right) Representative immunostainings for IBA1 in brain sections at day 21 in indicated groups. Scale bars, 100 μm. (Left) Histological percentage of IBA-1+ among total tissue in brain at 0-, 8- and 21 days post-infection. ns, not significant; *p ≤ 0.05; **p ≤ 0.01; ***p ≤ 0.001; (two-tailed unpaired t-test for C; two-way ANOVA followed by Sidak’s multiple comparisons test for D; two-tailed unpaired t-test for E; two-way ANOVA followed by Fisher’s LSD multiple comparisons test for F-G). Data represent the pool of 2 independent experiments (B, C, D, E) or are representative of at least 2 independent experiments (F, G). Bars and horizontal lines represent mean ± SEM.

Next, we determined the role of oligodendrocyte-derived IL-33 in CD8+ T cell-mediated CNS autoimmune disease. To this end, we crossed IL-33 cKO mice to our previously described MOG-GP line (Page et al., 2019). MOG-GP mice express the lymphocytic choriomeningitis virus (LCMV) glycoprotein (LCMV-GP) as a neo-self-antigen specifically in oligodendrocytes. For our study, we compared mice that were either sufficient (MOG-GP) or deficient for oligodendrocyte-derived IL-33 (IL-33 cKO MOG-GP). To initiate an LCMV-GP-neo-self-specific CD8+ T cell response we infected mice intravenously (i.v.) with an attenuated variant of LCMV, referred to as “rLCMV-GP33” (Page et al., 2019). In this variant, the LCMV glycoprotein (GP) ectodomain has been replaced by the vesicular stomatitis virus glycoprotein (VSVG) ectodomain, retaining the LCMV-GP leader sequence, which comprises the H-2D^b^-restricted immunodominant GP_33-41_ epitope (Page et al., 2021). MOG-GP mice are not immunologically tolerant to LCMV-GP expressed in their oligodendrocytes. As a result, rLCMV-GP33 infection elicits a polyclonal endogenous CD8+ T cell response, which attacks LCMV-GP-expressing oligodendrocytes, causing oligodendrocyte loss and demyelination (Page et al., 2019). Following rLCMV-GP33 infection, IL-33 cKO MOG-GP mice exhibited a significantly milder CNS disease course than IL-33-competent MOG-GP controls (Fig. 1D). WT mice without MOG-GP transgene remained healthy during the entire three-week observation period, as expected (Fig. 1D). At the peak of disease on day 10, the number of Olig2+ oligodendrocytes was clearly reduced in the brain and spinal cord of MOG-GP controls but in IL-33 cKO MOG-GP mice was at least partially preserved (Fig. 1E). At a later stage of disease (day 21), immunohistochemical staining against amyloid precursor protein (APP) revealed fewer injured axonal spheroids in IL-33 cKO MOG-GP mice than in MOG-GP mice (Fig. 1F). In addition, IL-33 cKO MOG-GP showed a reduction in the tissue area made up by activated IBA-1^+^ microglia cells (Fig. 1G) and GFAP+ astrocytes (Supp. Fig. 1A-B), respectively, both on day 10 and day 21.

Overall, these results suggested that oligodendrocyte-derived IL-33 enhances the severity of CNS autoimmune disease and associated immunopathology, as evidenced by increased axonal damage, phagocyte activation, and astrogliosis.

### IL-33 promotes the persistence of self-reactive CD8+ T cells in the CNS by driving their effector differentiation from TCF-1^high^ stem-like precursors

Next, we performed flow cytometry and immunohistochemistry to enumerate CNS-infiltrating T cells in MOG-GP and IL-33 cKO MOG-GP mice at early (day 8) and later (day 21) stages of the disease (Fig. 2A-C, Supp. Fig. 1-A, C, and D). By flow cytometric analysis on day 8, both the total and GP_33-41_-specific CD8+ T-cell counts were comparable in the two groups (Fig. 2B-C). However, by day 21, the GP_33-41_-specific CD8+ T cells as well as the overall counts of CD8+ and CD4+ T cells, were significantly lower in the brains of IL-33 cKO MOG-GP mice (Fig. 2A-C and Supp. Fig. 1-A, D). Similarly, a histological assessment of CNS infiltrating CD8+ T cells revealed a reduced density of CD8+ T cells evident from day 10 onwards (Supp. Fig. 1A, C). CD4+ T cell infiltrates were also detected in the inflamed CNS, but at approximately ten times lower density than CD8+ T cells (Supp. Fig. 1A, C-D). At a late time point (day 21) the density of CD4+ T cells was significantly lower in IL-33 cKO MOG-GP mice than in MOG-GP mice. In addition, only very few B cells were observed in tissues, but they were not statistically differently abundant in the CNS of the two genotypes (Supp. Fig. 1A, E). In contrast to the above findings relating to CD8+ T cells in the CNS, splenic GP_33-41_-specific CD8+ T cell counts in the two genotypes of mice were not significantly different at either time point (Fig. 2D). This suggests that the reduction of CNS-infiltrating CD8+ T cells in IL-33 cKO MOG-GP mice was not a result of impaired initial expansion within secondary lymphoid organs.

**Figure 2:**
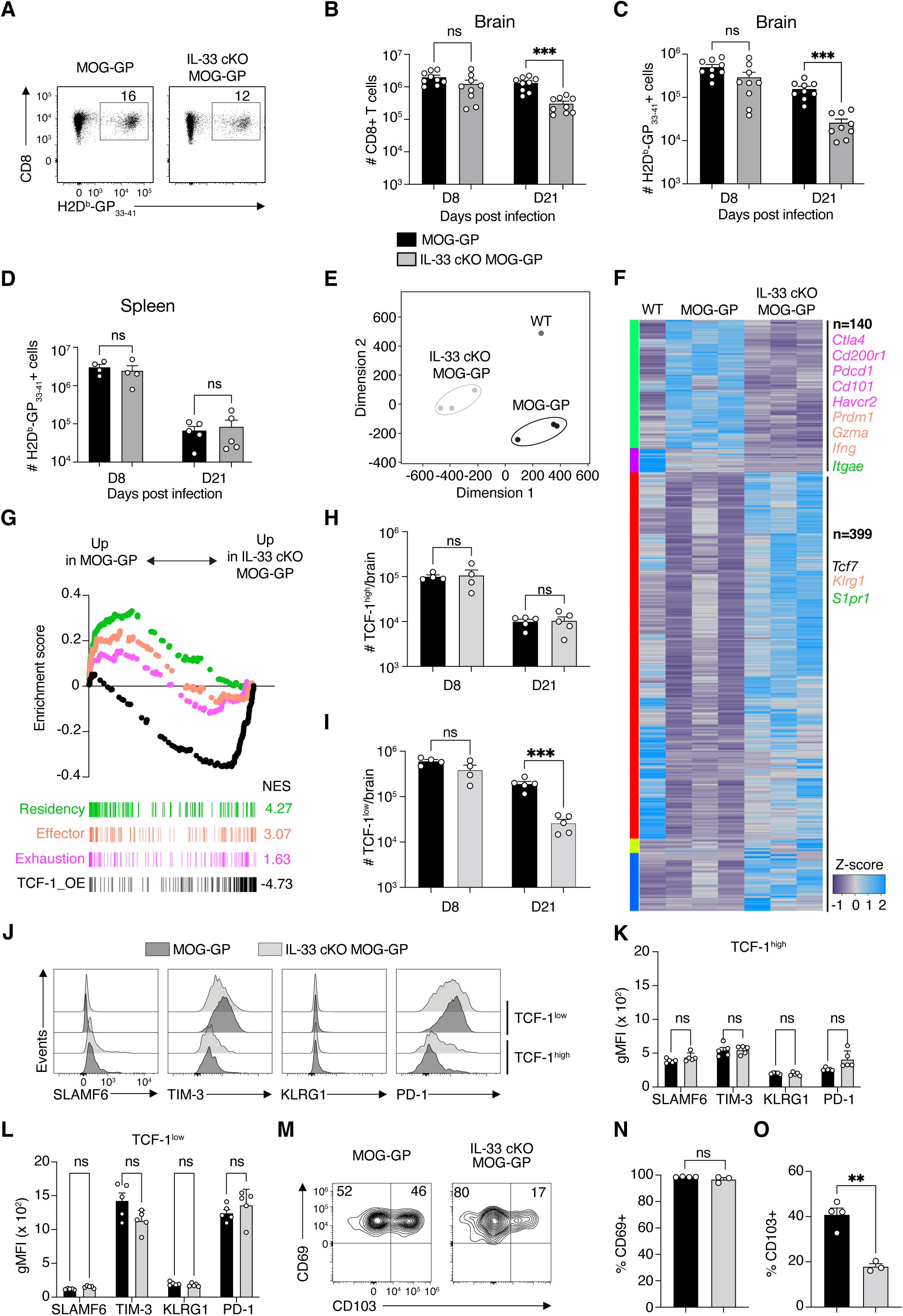
IL-33 promotes the persistence of self-reactive CD8+ T cells in the CNS by driving their effector differentiation from TCF-1^high^ stem-like precursors. **(A-D, H-O)** MOG-GP and IL-33 cKO MOG-GP mice were infected i.v. with 10^4^ PFU rLCMV-GP33. **(A)** Representative flow cytometry dot plot of brain-infiltrating H-2D^b^-GP_33-41_-specific CD8+ T cells at 8 days post-infection. Numbers over the gate indicate the frequency of H-2D^b^-GP_33-41_-specific CD8+ T cells. **(B)** Flow cytometric enumeration of brain-infiltrating and (**C**) H-2D^b^-GP_33-41_-specific CD8+ T cells at 8- and 21-days post-infection. **(D)** Flow cytometric enumeration of splenic H-2D^b^-GP_33-41_-specific CD8+ T at 8- and 21-days post-infection. **(E-G)** RNA-seq of FACS-sorted brain-infiltrating H-2D^b^-GP_33-41_-specific CD8+ T cells from WT, MOG-GP, and IL-33 cKO MOG-GP mice at 21 days post-infection with 10^4^ PFU rLCMV-GP33. **(E)** Multidimensional scaling (MDS) plot FACS-sorted H-2Db-GP_33-41_-specific CD8+ T cells in indicated groups. **(F)** Heatmap of Z-score of differentially expressed genes (DEGs) (FC ζ 1.5; FDR ≤ 0.05) in MOG-GP versus IL-33 cKO MOG-GP (n = 3 mice/group). The corresponding RNA expression of the selected DEGs obtained in WT brain is indicated in the left column of the heatmap (n = 1 pool of 4 mice). **(G)** GSEA of a signature of residency, exhaustion, effector differentiation, and TCF-1 OE in a ranked list of genes differentially expressed by brain-infiltrating H-2D^b^-GP_33-41_-specific CD8+ T cells from MOG-GP versus IL-33 cKO MOG-GP mice. NES: normalized enrichment score. **(H)** Absolute quantification of TCF-1^high^ and **(I)** TCF-1^low^ H-2D^b^-GP_33-41_-specific CD8+ T cells at 8- and 21-days post-infection. **(J)** Representative flow cytometry histograms of the indicated protein expression in brain-infiltrating TCF-1^high^ and TCF-1^low^ H-2D^b^-GP_33-41_-specific CD8+ T cells at 21 days post-infection. **(K)** gMFI of SLAMF6, TIM-3, KLRG1 and PD-1 in TCF-1^high^ and **(L)** TCF-1^low^ H-2D^b^-GP_33-41_-specific CD8+ T cells at 8- and 21-days post-infection. **(M)** Representative flow cytometry dot plot of CD69 and CD103 expression in brain-infiltrating H-2D^b^-GP_33-41_-specific CD8+ T cells at 21 days post-infection. **(N)** Frequencies of CD69+ and **(O)** CD103+ H-2D^b^-GP_33-41_-specific CD8+ T cells at 21 days post-infection.ns, not significant; *p σ 0.05; **p σ 0.01; ***p σ 0.001 (unpaired t-test for B-D, H-I, N-O. unpaired t-test with Benjamini, Krieger and Yekutieli multiple comparisons test for K-L ). Data represent the pool of 2 independent experiments (B-D), have been performed once (E-G), or are representative of at least 2 independent experiments (H-O). Bars and horizontal lines represent mean ± SEM.

Further, we infected mice intracranially (i.c.) with rLCMV-GP33 to test the effects of oligodendrocyte-derived IL-33 on CD8+ T cells in the context of transient viral infection of the CNS. Despite viral clearance and mild meningoencephalitis in wild-type (WT) animals (Pinschewer et al., 2010) infected with attenuated LCMV variants, such infections establish a pool of antiviral brain resident memory T cells (TRM) (Steinbach et al., 2016). Intriguingly, both IL-33 cKO and WT mice showed similar numbers of persistent brain-resident GP_33-41_-specific memory CD8+ T cells (Supp. Fig. 2A-B). Also, the expression levels of tissue-resident memory markers CD69 and CD103 on GP_33-41_-specific CD8+ T cells (Supp. Fig. 2C-D), the cytokine production by these cells upon peptide re-stimulation (Supp. Fig. 2E-G), and their expression of TCF-1 and of the inhibitory receptor PD-1 were similar in both groups of mice (Supp. Fig. 2H-J). This suggested that, unlike in autoimmunity, oligodendrocyte-derived IL-33 does not appear to play a critical role in the generation and persistence of antiviral CD8 T cells following transient viral CNS infection.

Next, we explored how oligodendrocyte-derived IL-33 alters the transcriptional landscape of self-reactive CD8+ T cells in the inflamed CNS. To this end, we performed RNA-seq on FACS-sorted GP_33-41_-specific CD8+ T cells from the brain of WT, MOG-GP, and IL-33 cKO MOG-GP mice, on day 21 after rLCMV-GP33 infection. Multidimensional scaling (MDS) analysis of differentially expressed genes (DEGs) segregated GP-reactive CD8+ T cells into three distinct clusters corresponding to the three different mouse genotypes (Fig. 2E). A comparison of GP_33-41_-specific CNS-infiltrating CD8+ T cells from MOG-GP and IL-33 cKO MOG-GP mice identified 539 DEGs (FC ≥ 1.5; FDR < 0.05; Fig. 2F). In the absence of oligodendrocyte-derived IL-33 we observed reduced expression of *Itgae* encoding for the retention molecule CD103 and of several genes encoding for inhibitory receptors including *Pdcd1*, *Havcr2*, *Cd200r1*, *Ctla4*, and *Cd101*. These inhibitory receptors indicate recent antigen contact and in the context of persistent viral infection identify cells in a state termed T cell exhaustion (Fuertes Marraco et al., 2015). Consistent with these observations, gene set enrichment analysis (GSEA) corroborated that CNS-infiltrating GP_33-41_-specific CD8+ T cells from IL-33 cKO MOG-GP mice exhibited a reduction in gene expression signatures associated with tissue residency and effector/exhaustion gene expression (Fig. 2F). Furthermore, analysis of DEGs revealed a relative enrichment of *Tcf7* transcripts encoding for the stemness-related TF TCF-1 (Fig. 2F). We thus interrogated on a broader scale whether known TCF1-associated genes were enriched in the absence of oligodendrocyte-derived IL-33 (Fig. 2G). GSEA showed that GP_33-41_-specific CD8+ T cells in the brain of IL-33 cKO MOG-GP mice exhibited a more pronounced TCF-1-associated gene expression signature as established in the context of chronic viral infection and cancer (Fig. 2G, (Wu et al., 2016)). Altogether, these findings demonstrated that oligodendrocyte-derived IL-33 plays a critical role in shaping the transcriptional landscape of CD8+ T cells in the context of CNS autoimmunity. These effects induced the regulation of effector, exhaustion, and residency signatures, as well as TCF-1-regulated genes.

We further investigated how oligodendrocyte IL-33 impacts the expression of TCF-1 and of inhibitory receptors by GP_33-41_-specific CNS-infiltrating CD8+ T cells. At both early (day 8) and late (day 21) stages of CNS autoimmunity, GP_33-41_-specific CD8+ cells in the brain of IL-33 cKO MOG-GP mice contained a higher percentage of TCF-1^high^ cells than those in MOG-GP controls (Supp. Fig. 3A-B). This proportional increase in TCF-1^high^ cells amongst autoreactive CNS-infiltrating CD8+ T cells was confined to the CNS of IL-33 cKO MOG-GP mice, whereas frequencies and absolute numbers of TCF-1^high^ GP_33-41_-specific CD8+ T cells in the spleen were similar to MOG-GP controls (Supp. Fig. 3C-D). Notably also, absolute numbers of TCF-1^high^ brain-infiltrating CD8+ T cells were comparable in the two experimental groups (Fig. 2H), both on day 8 and day 21, indicating that oligodendrocyte IL-33 is not required for the recruitment and retention of TCF-1^high^ self-reactive CD8+ T cells in the autoimmune CNS. Conversely, the pool of TCF-1^low^ effector-differentiated GP_33-41_-specific CD8+ T cells in the brain of IL-33 cKO MOG-GP mice started at a level comparable to MOG-GP control mice on day 8 but contracted to ∼10-fold lower levels by day 21 (Fig. 2I). These findings were best compatible with an impaired differentiation of self-reactive effector CD8+ T cells from TCF-1^high^ precursors locally in the brain when oligodendrocyte-derived IL-33 was lacking.

We next examined if oligodendrocyte-derived IL-33 affects the expression of the inhibitory receptors SLAMF6, PD-1, TIM-3, and KLRG1 by self-reactive GP_33-41_-specific CD8+ T cells. By separately analyzing the TCF-1^high^ and TCF-1^low^ subsets of GP_33-41_-specific CD8+ T cells in the brain on day 21 after rLCMV-GP33 infection, we observed that the absence of IL-33 did not impact inhibitory receptor expression in either subset (Fig. 2J-L). Moreover, in agreement with the inhibitory function of TCF-1 on CD103 expression (Wu et al., 2020) and in keeping with an increased proportion of TCF-1^high^ brain-infiltrating CD8+ T cells in IL-33 cKO MOG-GP mice, we observed that self-reactive CD8+ T cells persisting in the brain of these animals expressed lower levels of CD103 while the CD69 levels were comparable to MOG-GP controls (Fig. 2M-O). Considering the established precursor–product relationship of TCF-1^high^ and TCF-1^low^ CD8+ T cell subsets, our findings suggest that oligodendrocyte-derived IL-33 plays a crucial role in sustaining and/or expanding the self-reactive CD8+ effector T cell pool in the brain, likely by promoting the differentiation of TCF-1^high^ stem-like cells into TCF-1^low^ effector cells.

### Therapeutic blockade of IL-33 signaling in the CNS mitigates the severity of CNS autoimmune disease

We finally explored whether IL-33 represents a druggable target in CD8+ T cell-driven autoimmune disease of the CNS and, hence, may be harnessed for therapeutic purposes. For this, we exploited an adeno-associated viral (AAV) vector expressing an IL-33 decoy receptor (AAV-ST2-Fc), which can be administered to mice as somatic gene therapy and blocks IL-33 signaling to CD8+ T cells (Kastner et al., 2024). We administered AAV-ST2-Fc intracranially (i.c.) to MOG-GP mice on day 8 after rLCMV-GP33 infection, thus around the time of disease onset when CD8+ T cell expansion and substantial infiltration has already taken place (see Fig. 2B). When compared to controls receiving an AAV control vector expressing irrelevant cargo (AAV-control), interventional AAV-ST2-Fc therapy significantly ameliorated disease progression from around day 14 onwards (Fig. 3A). We confirmed expression of the ST2-Fc decoy receptor in the serum and brain of intracranially injected mice on day 21 (Fig. 3B-C). As assessed histologically, spinal cord-infiltrating CD8+ T cells on day 21 were significantly reduced in AAV-ST2-Fc-treated mice (Fig. 3D-E), phenocopying to some extent the reduced T cell infiltrates in the brain of IL-33 cKO MOG-GP lacking IL-33 in oligodendrocytes (see Fig. 2B). Importantly also, the number of damaged axonal spheroids as visualized by immunohistochemical staining for APP was significantly lower in the AAV-ST2-Fc-treated group than in AAV-control-treated mice (Fig. 3F-G). Taken together, these data demonstrate IL-33 as a druggable tissue factor in the CNS, opening new avenues to mitigate the long-term progression and outcome of T cell-driven CNS autoimmune disease.

**Figure 3:**
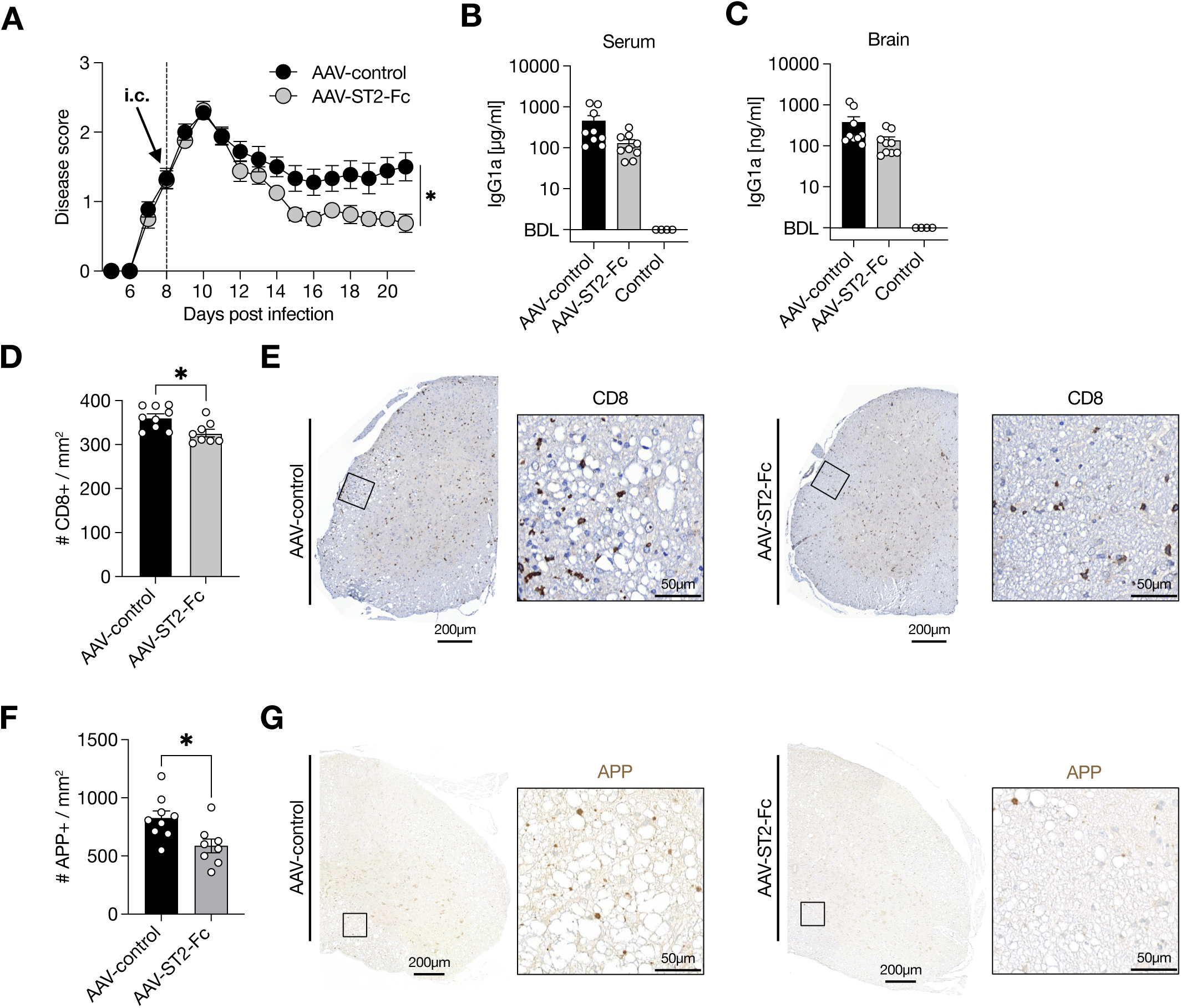
Local blockade of IL-33 pathway mitigates the severity of CNS autoimmune disease. MOG-GP mice were challenged i.v. with 10^4^ PFU rLCMV-GP33 and treated 8 days post-infection with AAV-ST2-Fc or the AAV8-Vl10 (control) by i.c. or i.m. **(A)** EAE disease course of i.c. post-challenge treatment (n = 9 MOG-GP mice treated with AAV-ST2-Fc; n = 8 MOG-GP mice injected with AAV8-Vl10 (control)), clinical scores are expressed as mean ± SEM. **(B)** Quantification of ST2-Fc and Vl10 control construct in the serum at day 21 post-infection. **(C)** Quantification of ST2-Fc and Vl10 control construct in the brain at day 21 post-infection. **(D)** Enumeration of CD8+ T cell infiltration in spinal cord tissues after AAV i.c. (left) and i.m. (right) treatment 21 days post-infection. **(E)** Representative immunostainings CD8 in spinal cord tissue 21 days post-infection **(F)** Enumeration of APP (Up) and Olig2+ (Down) per mm^2^ in spinal cord tissues after AAV i.c. (left) and i.m. (right) treatment 21 days post-infection. **(G)** Representative immunostainings APP in spinal cord tissue 21 days post-infection ns, not significant; *p ≤ 0.05 (two-way ANOVA followed by Sidak’s multiple comparisons test for A; two-tailed unpaired t-test for B-C; unpaired t-test for D, F). Bars and horizontal lines represent mean ± SEM.

## Discussion

Our study provides evidence that oligodendrocyte-derived IL-33 plays an important role in shaping the differentiation of self-reactive CD8+ T cells. This has implications for the disease progression and tissue destruction in a model of CNS autoimmunity. Absence of IL-33 in oligodendrocytes resulted in reduced persistence of self-reactive CD8+ T cells in the CNS, alongside with diminished transcriptional signatures of tissue residency and effector cell differentiation in brain-infiltrating autoreactive CD8+ T cells. Likewise, in the absence of IL-33 in oligodendrocytes, TCF-1^High^ stem-like CD8+ T cells exhibited an impaired capacity to replenish the effector CD8+ T cell pool necessary to sustain long-term autoimmune responses.

We previously demonstrated that self-reactive CD8+ T cells in the CNS undergo a distinct differentiation trajectory, characterized by the acquisition of an exhausted-like phenotype (Page et al., 2021). Consistent with a recent study (Grebinoski et al., 2022), our findings suggest that IL-33 regulates the stemness activity of autoreactive CD8+ T cells by which the pool of effector self-reactive CD8+ T cells are replenished and thus relies on stem-like progenitors in the affected tissue. This contrasts with other reports in a mouse diabetes model, which suggests that the pathogenic effector CD8+ T cell population primarily originates from a CD8+ T cell population located away from the antigen source in the peripheral lymph node (Abdelsamed et al., 2020; Gearty et al., 2022). This discrepancy could be due to differences in the tissue microenvironments, especially for the CNS, where drainage of non-sequestered antigens to secondary lymphoid organs may be limited compared to the pancreas. As a result, compartmentalized immune responses may prevail as the disease progresses towards chronicity (Matejuk et al., 2021; Forrester et al., 2018).

Our study is in line with previous observations (Casey et al., 2012) and extends their relevance to CNS autoimmunity, emphasizing the critical role of IL-33 in promoting the differentiation of T cells into T_RM_. It has been shown that TCF-1 inhibits the tissue residency program of memory CD8+ T cells, mainly by repressing the expression of the retention molecule CD103 in the lung (Wu et al., 2020). Our observations suggest that in the absence of IL-33 in oligodendrocytes, autoreactive CD8+ T cells remain preferentially trapped in a TCF-1^high^ stem-like cell stage, resulting in a compromised tissue-resident core signature marked by reduced CD103 expression. This appears to be particularly relevant in the context of chronic autoimmune conditions, where autoreactive T cells likely encounter antigens over a prolonged periods of time. Of note, this effect on CD8+ T cells was confined to the CNS, as IL-33 deficiency in oligodendrocytes did not result in redistribution or an altered numbers of memory T cells in the spleen. Furthermore, our data also indicate that the role of oligodendrocyte-derived IL-33 in T_RM_ generation and maintenance following transient viral infection, and thus transient antigen exposure to T cells, is possibly compensated by other cellular sources of IL-33 within the CNS.

Our study furthermore suggests that IL-33 deletion in oligodendrocytes restrains clonal expansion of self-reactive CD8+ T cells that are supposed to interact and destroy antigenic oligodendrocytes, leading to consecutive axonal alterations. As a result, IL-33-mediated effects could potentially promote the emergence of new T cell clones through antigen release and epitope spreading, thereby perpetuating inflammation and disease progression in demyelinating disorders like MS.

It is well established that IL-33 initiates signaling cascades during T cell activation, differentiation, and expansion in secondary lymphoid tissues (Guo et al., 2022, Brunner et al., 2024). The finding that oligodendrocyte-derived IL-33 in T cell fate bifurcation within the inflamed tissue bears implications for potential future therapeutic interventions, as our therapeutic approach suggested as well. Our finding of successful therapeutic intervention by AAV-based delivery of a IL33 decoy receptor suggests that targeting local IL-33 signaling might offer a therapeutic opportunity to diminish compartmentalized inflammation, which has been observed in more advanced disease stages of multiple sclerosis (Allan et al., 2016). Such an approach could specifically address the involvement of tissue-resident CD8+ T cells (Vincenti et al., 2022), which are considered to be more resistant to standard therapies that primarily aim to curtail the expansion and migration of circulating immune cells into the CNS. Beyond autoimmune diseases, IL-33 has been considered a valuable cytokine for enhancing anti-tumor immunotherapy (Gao et al., 2015, Kallert et al., 2017), as it fosters CD8+ T cell tumor infiltration. However, whether IL-33 improved infiltration acts through TCF-1 signaling in this latter context remains to be demonstrated. Additionally, therapeutic strategies aimed at providing alarmins to boost tumor infiltration and the anti-tumoral function of CD8+ T cells should also take into account that exogenous IL-33 may confer stem-like properties to cancer cells, potentially promoting carcinogenesis (Fang et al., 2017).

A prerequisite for the formation of effector cells during viral or bacterial infection is the downregulation of TCF-1 (Zhou et al., 2010). Growing evidence indicates that inflammatory cues such as IL-12 and type I IFN are acting as inhibitors of TCF-1 expression thus facilitating effector CD8+ differentiation (Danilo et al., 2018). Precisely, in a model of chronic LCMV infection, blockade of type I interferon, a driver of T cell exhaustion, promoted the expansion of TCF-1^high^ stem-like cells CD8+ T cells (Wu et al., 2016). Similarly, our data demonstrate that selective abrogation of IL-33 in oligodendrocytes leads to an increase of TCF-1^high^ cells in the CNS. Given the notion that IL-33 combined with IL-12 promotes IFN-γ production in CD8+ T cells (Yang et al., 2011), it is thus tempting to speculate that IL-33-mediated act synergistically with IL-12 to control TCF-1 expression and T cell effector differentiation.

In summary, our work in the context of CNS autoimmunity provides new insights into the role of oligodendrocyte-derived IL-33, which facilitates the differentiation of self-reactive TCF-1^high^ stem-like cells into terminally differentiated effector cells possessing high encephalitogenic potential. This insight into the molecular basis of autoreactive T cell persistence and expansion may thus contribute to defining novel immunotherapeutic strategies for chronic CNS autoimmune disorders.

## Methods

### Mice

C57BL/6J WT were obtained from Charles River (France). C57BL/6J MOG-GP (MOGi^Cre/+^: Stop-GP^flox/+^) mouse line was generated by crossing mice expressing the Cre-recombinase under the control of the oligodendrocyte-specific promoter (C57BL/6J MOGiCre (Hövelmeyer et al., 2005); with C57BL/6J Stop-GP mice (Page et al., 2019)). C57BL/6J IL-33 cKO MOG-GP mouse line was generated by crossing IL-33^flox/flox^ mice (Chen et al., 2015) with C57BL/6J MOG-GP mice. All mice were lodged under specific-pathogen-free P2 conditions in the animal facilities of the University Medical Center of Geneva. Male and female sex and age-matched mice between six weeks and twelve weeks of age were used for experiments. All animal experiments were authorized by the cantonal veterinary office of Geneva and performed in agreement with the Swiss law for animal protection.

### Virus Infection

Recombinant LCMV were generated and propagated on baby hamster kidney 21 cells BHK21 (ATCC® CCL-10) (Flatz et al., 2006). Virus stocks were titrated using MC57G cells (ATCC® CRL-2295) according to established methods (Kallert et al., 2017). The recombinant rLCMV-GP33 expresses the signal sequence of LCMV glycoprotein (GP) harboring the GP_33-41_ epitope fused to the VSV GP instead of the LCMV full-length GP. For transient virus infection in the brain, 10^4^ plaque-forming units (PFU) of rLCMV-GP33 were diluted in 30 μl of minimum essential medium (Gibco) before intracranial (i.c.) injection. For the peripheral challenge, mice were intravenously (i.v.) injected into the tail vein with 10^4^ PFU of rLCMV-GP33 in 100 ul. Infected animals were monitored daily for the occurrence of classical experimental autoimmune encephalomyelitis (EAE) symptoms and scored as follows. 1, flaccid tail; 2, impaired righting reflex and hind limb weakness; 3, complete hind limb paralysis; 4, complete hind limb paralysis with partial fore limb paralysis; 5, moribund. Severely diseased animals were immediately sacrificed.

To block IL-33 signaling, we used an AAV-ST2-Fc described by (Kastner et al., 2024). In short, the ST2-Fc construct is composed of a signal peptide and the extracellular domain of ST2 (NCBI, Genbank: M24843.1) coupled C-terminally to the mouse IgG1a constant domain. The ST2-Fc was cloned into the pENN.AAV.CB7.CI AAV expression plasmid, which contains a CAG-promoter-driven expression cassette flanked by AAV2 inverted terminal repeats (ITRs), obtained from the PENN Vector Core (Perelman School of Medicine, University of Pennsylvania, PA, USA) under MTA. As control, we used an AAV vector expressing the antibody VI10 (Kalinke et al., 1996; Ertuna et al., 2021). AAV vector production (in an AAV8 capsid format) and titration were performed by the Viral Vector Facility (VVF) of the Neuroscience Center Zurich (ZNZ), Switzerland. AAV vectors were administered to mice i.c. at a dose of 1 x 10^11^ viral genomes per mouse.

### Immunohistochemistry

Brain and spinal cord tissues were collected and fixed in 4% paraformaldehyde (PFA) overnight at 4°C. Dehydrated tissues were embedded in paraffin. Deparaffinized tissue sections (2 μm) were incubated with Dako REAL peroxidase-blocking solution (Dako, K0672) to inactivate endogenous peroxidases and unspecific binding was blocked (PBS/1% BSA). After antigen retrieval, sections were incubated with primary antibody for IL-33 (R&D Systems, Cat# AF3626) and Olig2 (IBL, Cat# 18953). Bound primary antibodies were visualized with anti-goat Alexa Fluor 488–labeled secondary antibodies or peroxidase-labeled secondary antibodies, followed by opal 570 signal amplification system (opal 570; Akoya, FP1488001KT). Nuclei were stained with DAPI (Invitrogen, D1306).

For bright-field microscopic images, bound primary antibodies (anti-CD8 (eBioscience, 14-0195-80); anti-CD4 (Cell signaling, #25229), anti-B220 (eBioscience, 14-0452-85, anti-GFAP (Dako, Z0334), anti-IBA-1(Wako, 019-1974), anti-APP (Millipore, MAB348) were visualized using peroxidase-coupled secondary antibody systems (Dako, Vector Laboratories) and polymerized 3,3′-diaminobenzidine (Dako). Nuclei were counterstained with hemalaun.

### Image analysis

Immunostained sections were scanned using Pannoramic Digital Slide Scanner 250 FLASH II (3DHISTECH) in 200× magnification. Automatic quantification of positive cells was performed using Visiopharm platform using a custom-made script. For representative images, the white balance was adjusted, and contrast was enhanced using the tools “levels,” “curves,” “brightness,” and “contrast” in Photoshop CS6 (Adobe). All modifications were acquired uniformly on the entire image.

### FACS analysis and sorting

The following fluorophore-conjugated antibodies for flow cytometry were purchased from Biolegend (anti-CD8a Pacific Blue (53-6.7), anti-CD8a Brillant violet 605 (53-6.7), anti-TIM-3 Brillant violet 605 (RMT3-23), anti-KLRG1 FITC (2F1), anti-PD-1 PerCP/Cyanine5.5 (29F.1A12), anti-CD69 FITC (H1.2F3), anti-CD103 Alexa Fluor 647 (2E7), anti-CD44 PerCP/Cyanine5.5 (IM7), anti-IFNg Brillant violet 421 (XMG1.2), anti-IL-2 PE (JES6-5H4), anti-TNF PE/Cyanine7 (MP6-XT22)), Cell signaling Technology (anti-TCF-1 Alexa Fluor 647 (C63D9)) and BD Biosciences (anti-SLAMF6 Brillant Ultraviolet 395 (13G3)). For detection of GP-specific CD8+ T cells, PE labelled Db-GP_33-41_-tetramer (provided by the National Institutes of Health Tetramer Core Facility) was used. Dead cells were excluded by staining with Zombie NIR dye (Biolegend). Most antibodies were used at a 1/100 dilution. Peripheral blood samples were obtained by facial vein puncture in heparin. Blood erythrocytes were lysed, and cells fixed using BD FACS Lysing Solution (BD Biosciences, 349202). For the preparation of CNS infiltrating leukocytes, mice were anesthetized and transcardially perfused with PBS. Brains were minced, and digested in DMEM with Collagenase A (1mg/ml, Roche) and DNaseI (0.1 mg/ml, Roche) for 1h at 37°C and homogenized using 70-μm cell strainers (BD). Leukocytes were separated using a discontinuous percoll gradient (30 / 70%). The remaining erythrocytes were lysed using RBC Lysis buffer (Biolegend,420301) for 3 min at RT. Surface staining was carried out with directly labeled antibodies in FACS buffer (2.5% FCS, 10 mM EDTA, 0.01% NaN3 in PBS). Isolated cells were quantified using AccuCheck Counting Beads (Invitrogen, PCB100). Intracellular staining of TCF-1 was performed using FoxP3/Transcription Factor Staining Buffer Set (eBioscience, 00-5523-00) according to manufacturer’s instructions. To assess intracellular cytokine production, isolated leukocytes were cultured for 5h in the presence of Monensin (BioLegend) and Brefeldin (BioLegend). Cells were stimulated with 1 μM KAVYNFATM peptide. Flow cytometric samples were acquired on a BD LSRFortessa (BD Biosciences) using BD FACSDiva (BD Biosciences, v8.0.2) and Attune NxT Acoustic Focusing Cytometer (Thermo Fisher, V3.1.2) using appropriate filter sets and compensation controls. Gates were assigned according to appropriate control populations. Data were analyzed using FlowJo software (V10).

### RNA-seq samples preparation

To perform RNA-seq, we FACS-sorted brain CD8 GP ^+^ T cells from WT, MOG-GP, and IL-33 cKO MOG-GP mice at 21 days after rLCMV-GP33 infection. Each biological replicate (n = 3 for MOG-GP, n = 3 for IL-33 cKO MOG-GP and n = 1 pool of 4 WT mice) represents 10^4^ sorted H-2D^b^-GP_33-41_-specific CD8+ T cells in RLT buffer with 1% ß-mercaptoethanol and processed with RNAeasy Micro Kit (Qiagen, 74004) following manufacturer’s instructions.

were sorted in RLT buffer with 1% ß-mercaptoethanol and processed with RNAeasy Micro Kit (Qiagen, 74004) following manufacturer’s instructions.

Quality and integrity of RNA were assessed with the fragment analyzer by using the standard sensitivity RNA Analysis Kit (Advanced Analytical, DNF-471). All samples selected for sequencing exhibited an RNA integrity number over 8. RNA-seq libraries were generated using 100 ng total RNA of a non-stranded RNA Seq, massively parallel mRNA sequencing approach from Illumina (TruSeq mRNA Library Preparation, Illumina, 20020594). Libraries preparation was performed on the Beckman Coulter’s Biomek FXP workstation. For accurate quantitation of cDNA libraries, the fluorometric-based system QuantiFluor™dsDNA System was used (Promega, E2670). The size of final cDNA libraries was determined by using the dsDNA 905 Reagent Kit (Advanced Bioanalytical, DNF-905) exhibiting a sizing of 300 bp on average. Libraries were pooled and sequenced on the Illumina HiSeq 4000 sequencer (SE; 1 x 50 bp; 30-35 Mio reads/sample). Basecalls generated by Illumina’s Real Time Analysis (RTA) software were demultiplexed to fastq files with bcl2fastq (v2.17.1.14). The quality check was done using FastQC (Andrews S. (2010). FastQC: a quality control tool for high throughput sequence data. Available online at: http://www.bioinformatics.babraham.ac.uk/projects/fastqc) (version 0.11.8, Babraham Bioinformatics).

### RNA-seq data processing

FastQ reads were mapped to the ENSEMBL reference genome (GRCm38.96) using STAR version 2.4.0j (Dobin et al., 2013) with standard settings except that any reads mapping to more than one location in the genome (ambiguous reads) were discarded (outFilterMultimapNmay=1).

A unique gene model was used to quantify the number of reads per gene. Briefly, the model considers all annotated exons of all annotated protein coding isoforms of a gene to create a unique gene where the genomic regions of all exons are considered coming from the same RNA molecule and merged together.

All reads overlapping the exons of each unique gene model were reported using featureCounts version 1.4.6-p1 (Quinlan and Hall, 2010). Gene expressions were reported as raw counts and in parallel normalized in RPKM to filter out genes with low expression values (1 RPKM) before calling for differentially expressed genes. Library size normalizations and differential gene expression calculations were performed using the package edgeR (v3.28.0) (Robinson et al., 2010) designed for the R software (v3.6.2). Only genes having a significant fold-change (Benjamini-Hochberg corrected p-value < 0.05) were considered for the rest of the RNA-seq analysis.

### Gene set enrichment analysis

Gene set enrichment analysis (GSEA) was performed with gene sets identified using published microarray data sets. Effector and exhaustion signature originated from (Doering et al., 2012) studying the transcriptome of distinct CD8+ T cell differentiation states after LCMV Armstrong and LCMV clone 13 infection. TCF-1-enriched gene signatures originated from microarray analysis of TCF-1-overexpressing CD8+ T cells (Wu et al., 2016). Residency signature originated from studying the role of RUNX3 on CD8+ T cell residency in non-lymphoid tissues and tumors (Milner et al., 2017).

### Statistical analysis

Statistical parameters including the exact value of n, the dispersion, the precision of measures and the statistical significance are reported in the figures and figure legends. In figures, asterisks denote statistical significance as calculated by Student’s t-test and two-way ANOVA with Šidák post-test (ns, not significant; *, p < 0.05; **, p < 0.01; ***, p < 0.001). Statistical analysis was performed in GraphPad Prism 9.

## Data availability

Data are available in the article and Supplementary data file or from the corresponding author upon reasonable request. The raw and processed RNA-seq data have been deposited to the Gene Expression Omnibus (GEO) under accession numbers GEO: GSExxxx.

## Acknowledgments

We thank the iGE3 Genomics Platform at the University of Geneva for RNA-sequencing sequencing, the Flow Cytometry core facility at the CMU of the University of Geneva for help with flow cytometry. D.M. was supported by the Swiss National Science Foundation (310030_173010) and the Swiss MS Society. We thank the NIH Tetramer Core Facility for providing the H2-Db_LCMV-GP33-41-PE tetramer.

## Author contributions

N.F. and N.P. performed experiments and corresponding data analysis. N.P. and D.D.P helped in designing experiments. S.L. performed sequencing data analysis. M.K. developed and applied computer-assisted image analysis algorithms. B.K., G.D.L., M.P., I.V., A.L.K. and Y.I.E. were involved in animal experimentations and mouse tissue collection. I.W. performed the histological tissue processing and staining. A.L.K., Y.I.E. and D.D.P. provided adenovirus-associated viral vectors. N.F., N.P. and D.M. wrote the manuscript. D.M. designed the project.

## Corresponding author

Correspondence to Doron Merkler.

---

**Supplementary Figure 1:**
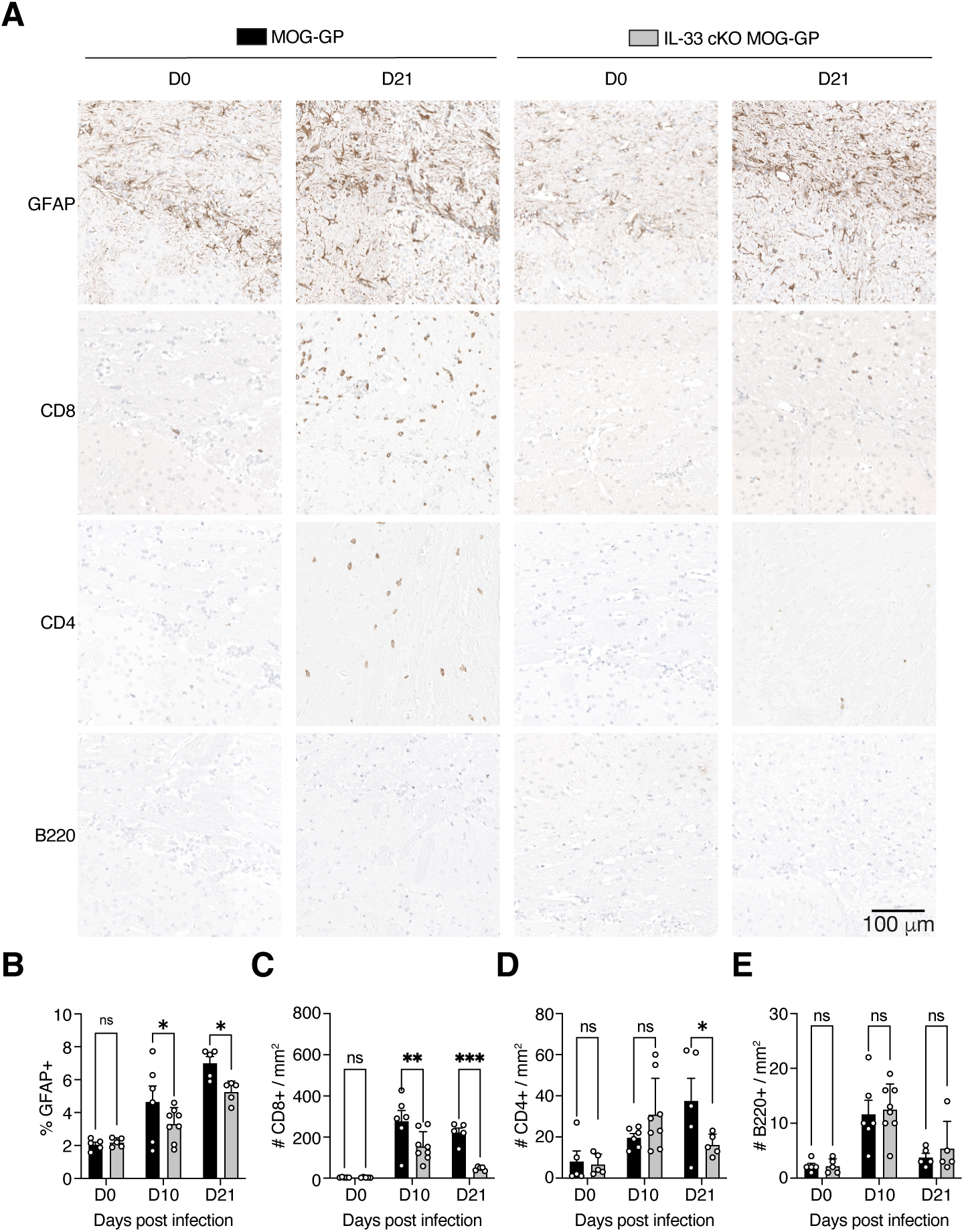
Absence of oligodendrocyte-derived IL-33 curtails brain inflammation and CD8+ T cells persistence. MOG-GP and IL-33 cKO MOG-GP mice were infected i.v. with 10^4^ PFU rLCMV-GP33. **(A)** Representative immunostainings for APP, IBA-1, GFAP, CD8, CD4, and B220 in brain sections at indicated time points in indicated groups. Scale bars, 100 μm. **(B)** Frequencies of GFAP+ area among total tissue area from brain sections at indicated time points in indicated groups. **(C)** Quantification of CD8+, **(D)** CD4+, and **(E)** B220+ cells per mm2 from brain sections at indicated time points in indicated groups. ns, not significant; *p ≤ 0.05; **p ≤ 0.01; ***p ≤ 0.001 (two-way ANOVA followed by Fisher’s LSD multiple comparisons test for B-E). Data represent the pool of 2 independent experiments (B-E). Bars and horizontal lines represent mean ± SEM.

**Supplementary Figure 2:**
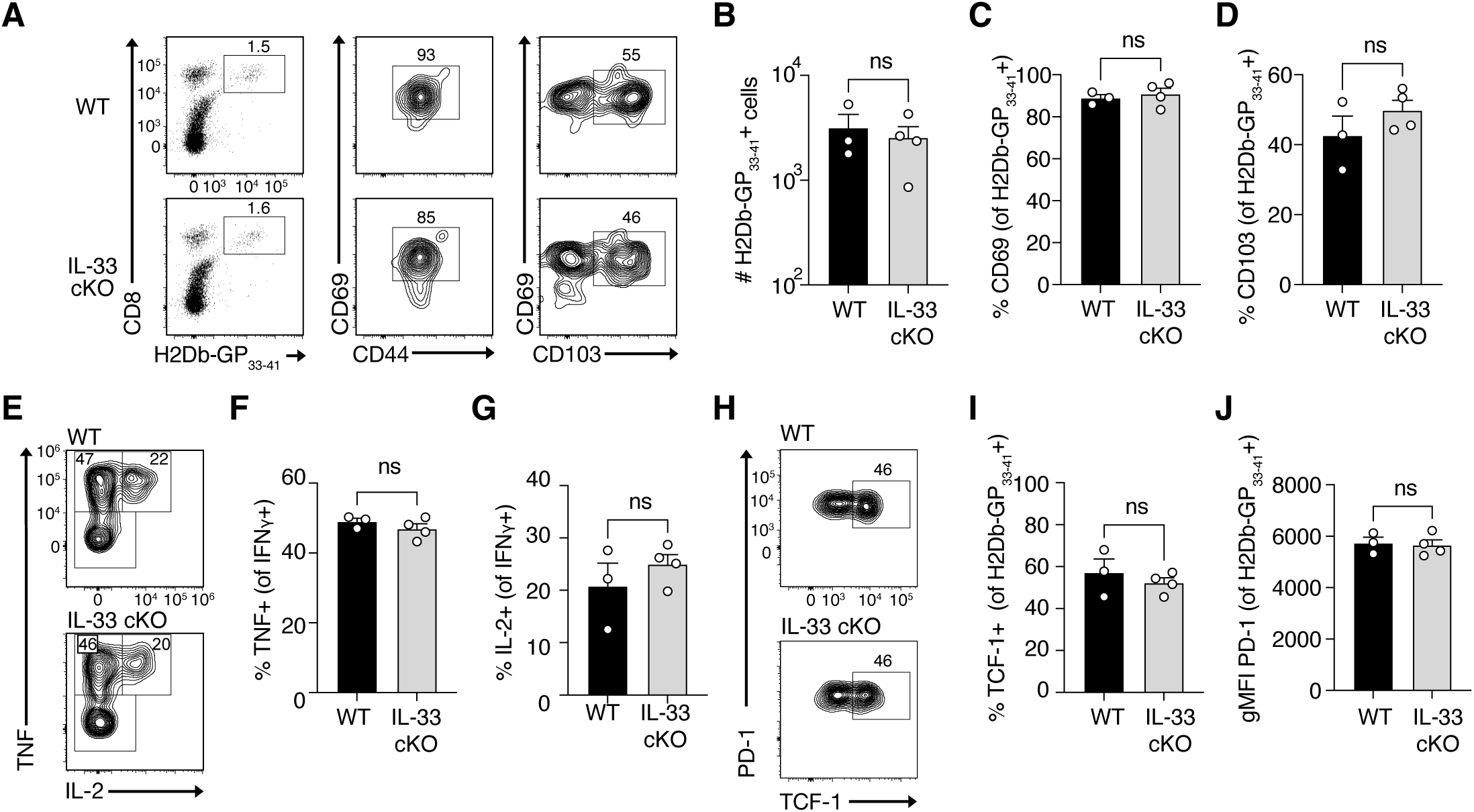
Oligodendrocyte-derived IL-33 is dispensable for classical brain T_RM_ formation and maintenance. WT and IL-33 cKO mice were challenged i.c. with 10^4^ PFU rLCMV-GP33 and brain-derived H-2D^b^-GP_33-41_-specific CD8+ T cells were isolated for FACS analysis at 6 weeks after infection. **(A)** Representative flow cytometry gating of H-2D^b^-GP_33-41_-specific CD8+ T cells and their respective CD69, CD103, and CD44 expression. Numbers indicate the frequency of positive cells. **(B)** Absolute counts of H-2D^b^-GP_33-41_-specific CD8+ T cells and percentages of **(C)** CD69+ and **(D)** CD103+ within the tetramer-specific population. **(E)** Representative flow cytometry plots of intracellular staining for TNF and IL-2 after *in vitro* stimulation with KAVYNFATC peptide. Cells were gated on IFNψ+ H-2D^b^-GP_33-41_-specific CD8+ T cells. Numbers indicate the frequency of cytokine producing cells within each gate. **(F)** Frequencies of TNF+ cells and **(G)** IL-2+ cells among IFNψ+ H-2D^b^-GP_33-41_-specific CD8+ T cells. **(H)** Representative flow cytometry plot of PD-1 and TCF-1 expression in H-2D^b^-GP_33-41_-specific CD8+ T cells. **(I)** Frequencies of TCF-1^high^ cells and **(J)** PD-1+ cells among H-2D^b^-GP_33-41_-specific CD8+ T cells. ns, not significant (two-tailed unpaired t-test for B-J). Data are representative of at least 2 independent experiments (A-J). Bars and horizontal lines represent mean ± SEM.

**Supplementary Figure 3:**
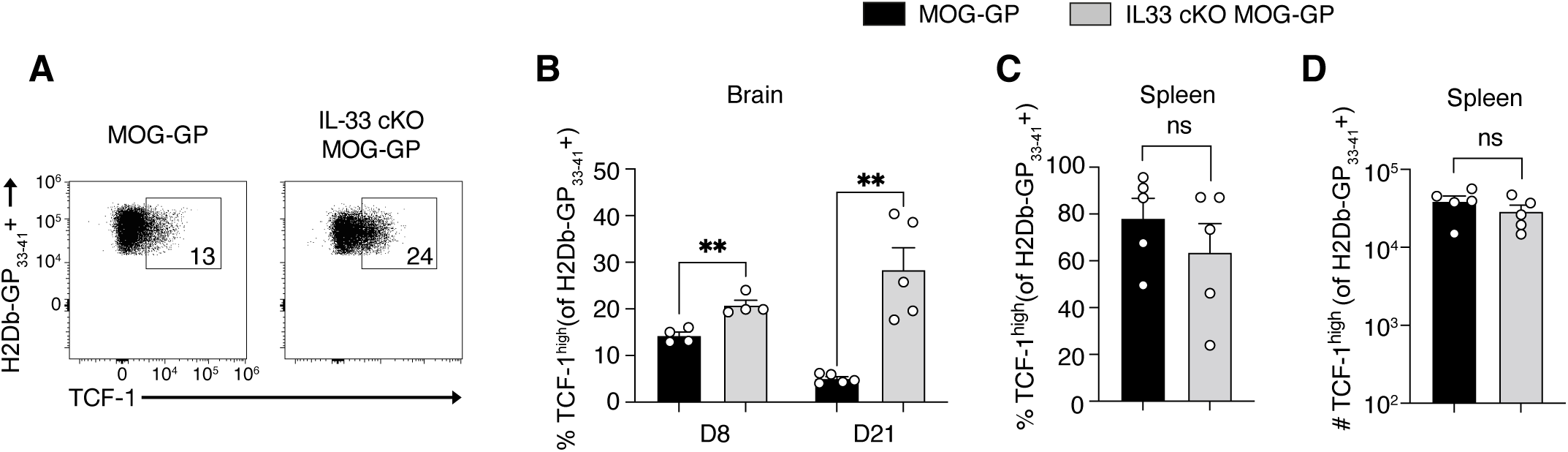
The proportion of TCF-1^high^ cells is higher in the brain and remains similar in the spleen in the absence of oligodendrocyte-derived IL-33. MOG-GP and IL-33 cKO MOG-GP mice were challenged i.v. with 10^4^ PFU rLCMV-GP33 and brain and splenic H-2D^b^-GP_33-41_-specific CD8+ T cells were analyzed at 8 days and 21 days post-infection. **(A)** Representative flow cytometry dot plot of TCF-1 expression in brain-infiltrating H-2D^b^-GP_33-41_-specific CD8+ T cells at 8 days post i.v. rLCMV-GP33 infection in MOG-GP and IL-33 cKO MOG-GP mice. **(B)** Percentages of TCF-1^high^ cells among brain infiltrating H-2D^b^-GP_33-41_-specific CD8+ T cells at 8- and 21-days post-infection. **(C)** Percentages and **(D)** absolute count of splenic TCF-1^high^ H-2D^b^-GP_33-41_-specific CD8+ T cells at 21 days post-infection. ns, not significant; **p ≤ 0.01 (two-tailed unpaired t-test for B-D). Data are representative of at least 2 independent experiments (A-D). Bars and horizontal lines represent mean ± SEM.

